# TARGETED MEMORY REACTIVATION DURING NON-RAPID EYE MOVEMENT SLEEP STRENGTHENS CONSOLIDATED DECLARATIVE MEMORIES

**DOI:** 10.64898/2026.08.24.746831

**Authors:** Malen D. Moyano, Lucila Capurro, María C. González, Luis I. Brusco, Cecilia Forcato

## Abstract

Targeted memory reactivation (TMR) during sleep can benefit recently acquired memories, but whether it can also influence memories after an initial period of consolidation remains unclear. Here, we tested whether auditory reactivation during non-rapid eye movement (NREM) sleep could strengthen declarative memories learned 24 h earlier. Twenty-six healthy young adults learned 30 sound–word associations and returned the following day for a 90-min nap. During NREM sleep, participants in the Reactivation group received incomplete reminders consisting of the learned sound followed by the first syllable of the associated word, whereas the No-Reactivation group slept under the same conditions without memory-related cues. Participants who received reminders showed significantly less forgetting, despite comparable training performance and sleep macroarchitecture. Across NREM sleep, reactivation was associated with greater slow oscillation and delta power, more slow oscillations and fast spindles, and greater slow oscillation–spindle co-occurrence. The memory benefit remained significant after adjusting for NREM physiological measures and in sensitivity analyses restricted to overlapping physiological ranges between groups. Cue-locked analyses revealed significant responses in the slow oscillation, delta, theta, and fast-spindle ranges, but the magnitude of these responses was not associated with memory change. These findings show that TMR during NREM sleep can benefit declarative memories after a 24-h consolidation interval and suggest that its effects extend beyond the immediate post-learning sleep period.

## Introduction

Sleep contributes to the stabilization of newly acquired memories (Diekelmann & Born, 2010; Rasch & Born, 2013; Lutz & Born, 2026). During non-rapid eye movement (NREM) sleep, neural representations formed during wakefulness are spontaneously reactivated, a process thought to support their long-term retention and integration into existing memory networks (Wilson & McNaughton, 1994; Ji & Wilson, 2007; Diekelmann & Born, 2010; Rasch & Born, 2013). This endogenous reactivation can be experimentally biased using targeted memory reactivation (TMR), in which sensory cues associated with prior learning are presented again during sleep (Rasch et al., 2007; Rudoy et al., 2009; Oudiette & Paller, 2013). Across different declarative memory tasks, cueing during NREM sleep has been shown to improve subsequent retention (Rasch et al., 2007; Rudoy et al., 2009; Schreiner & Rasch, 2015; Cairney et al., 2018).

Most TMR studies, however, present memory cues during the first sleep period after learning, often only a few hours after encoding (Rasch et al., 2007; Rudoy et al., 2009; Diekelmann et al., 2011; Antony et al., 2012; Cairney et al., 2018). At that point, the memory is still undergoing post-learning consolidation. It is therefore difficult to know whether TMR acts mainly by facilitating this ongoing process or whether sleep reactivation can also modify a memory after it has already passed through an initial consolidation period.

Studies of successive memory reactivation during wakefulness have shown that consolidated memories can be strengthened through repeated presentation of incomplete reminders (Forcato et al., 2007, 2009, 2011). By providing only part of the learned information, these reminders create a mismatch between what is expected and what is actually presented, generating a prediction error that can promote memory updating and strengthening (Fernández et al., 2016; Sinclair & Barense, 2018).

A similar effect of reminder structure has been observed during sleep. Using a sound-word association task, Forcato et al. (2020) compared incomplete reminders, consisting of the learned sound followed by the first syllable of the associated word, with complete reminders containing the full word. Both reminder types benefited memory when tested shortly after sleep, but only incomplete reminders produced a longer-lasting effect. These findings suggest that the effects of sleep reactivation depend on the information provided by the reminder.

Whether this principle also applies to memories reactivated after an initial consolidation interval is largely unknown. In mice, Rolls et al. (2013) re-exposed animals to an odor cue during sleep 24 h after fear conditioning and observed enhanced subsequent memory expression. This effect depended on protein synthesis in the basolateral amygdala, indicating that reactivation during sleep can engage plasticity mechanisms even after the initial post-learning period.

Comparable evidence for declarative memory in humans is still lacking.

Here, we tested whether TMR during NREM sleep can influence a declarative memory learned 24 h earlier. Participants learned sound-word associations and returned the following day for a daytime nap. During NREM sleep, one group received incomplete auditory reminders previously shown to produce persistent memory benefits, whereas a control group slept without memory-related cues. We predicted that reactivation would reduce forgetting relative to the control condition, providing evidence that the effects of sleep cueing are not limited to the first post-learning sleep period.

## Methods

### Participants

Fifty healthy young adults were recruited through advertisements distributed through social media. All participants provided written informed consent. The study was approved by the Biomedical Research Ethics Committee of the Instituto Alberto C. Taquini and the Human Ethics Committee of the Faculty of Medicine, University of Buenos Aires, and was conducted in accordance with the Declaration of Helsinki.

Because the protocol involved a daytime nap, when homeostatic sleep pressure is relatively low, stable NREM sleep and sufficient exposure to the reactivation protocol could not be achieved in all participants. Twenty-four participants were excluded because of technical problems during polysomnography (n = 5), failure to fall asleep (n = 4), less than 40 min of NREM sleep (n = 8), prolonged wakefulness during the reactivation period (n = 1), absence of slow-wave sleep or less than 2 min of slow-wave sleep (n = 2), or insufficient exposure to the reactivation protocol (n = 4). For the latter criterion, participants who received only 8 or 9 reminders were excluded, as this corresponded to reactivation of less than one third of the 30 learned associations. Participants who received four or fewer reminders were considered to have had negligible exposure to the reactivation protocol and were included in the No-Reactivation group. The final sample comprised 26 participants (mean age = 23.4 ± 0.9 years; 20 women).

Participants reported regular sleep–wake schedules and no history of neurological, psychiatric, or sleep disorders, psychoactive medication use, or night-shift work. They were asked to maintain their usual sleep schedule and to abstain from caffeine and alcohol on the experimental day.

### Experimental design

Participants were randomly assigned to a Reactivation group (R; *n* = 13) or a No-Reactivation group (NR; *n* = 13). The experiment was conducted over two consecutive days (Figure 1). On Day 1, participants arrived at the laboratory between 15:00 and 17:00 h and learned a set of sound-word associations. Participants returned approximately 24 h later and, after polysomnographic preparation, were given a 90-min daytime nap opportunity. Continuous white noise was presented through in-ear headphones in both groups. During NREM sleep, participants in the Reactivation group additionally received memory-related auditory cues, whereas the No-Reactivation group received no memory cues. After awakening, participants remained awake for 30 min before completing the memory test (Figure 1A), to reduce possible effects of sleep inertia (Tassi & Muzet, 2000). All participants had previously completed an adaptation nap in the laboratory.

**Figure 1.**
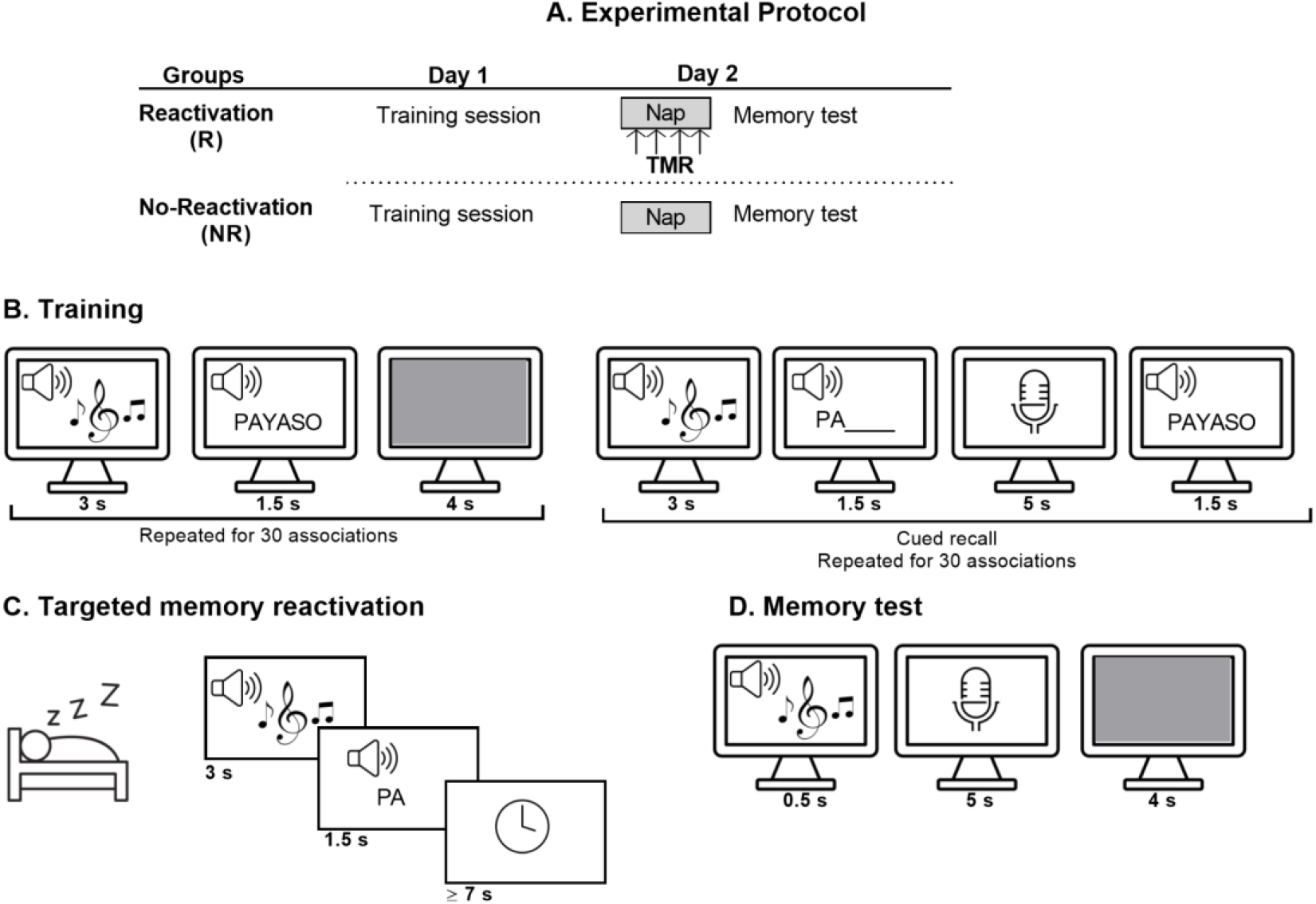
Experimental design and memory task. (A) Experimental protocol. On Day 1, participants learned 30 sound–word associations during a training session. Approximately 24 h later, they returned to the laboratory for a 90-min daytime nap with polysomnographic recording. During NREM sleep, participants in the Reactivation group received auditory memory cues, whereas participants in the No-Reactivation group slept under the same conditions without memory-related cues. After awakening, participants remained awake for 30 min and then completed the memory test. (B) Training. Each association was initially presented as an environmental sound followed by the corresponding three-syllable word in spoken and written form. Training performance was then assessed with cued recall: participants heard the environmental sound followed by the first syllable of the associated word and were asked to verbally report the complete word. Corrective feedback was provided after each response. (C) Targeted memory reactivation. During NREM sleep, participants in the Reactivation group received incomplete auditory reminders consisting of the learned environmental sound followed by the first syllable of the associated word. Reminders were delivered during stable NREM sleep and were separated by at least 7 s. (D) Memory test. After the nap, memory was tested by presenting each environmental sound alone. Participants verbally reported the associated word. No syllabic cue or corrective feedback was provided.

### Memory task

Participants learned 30 associations between environmental sounds and semantically related three-syllable Spanish words, using a task adapted from our previous studies (Forcato et al., 2020; Tassone et al., 2020). Words were presented through headphones using a prerecorded female voice.

### Training

Each association was first presented as an environmental sound (∼3 s), followed by the corresponding word presented simultaneously in spoken and written form (1.5 s). After presentation of all 30 pairs, memory was assessed by cued recall. Participants heard the sound followed by the first syllable of the associated word and were asked to verbally report the complete word. The correct word was then presented visually and auditorily as feedback (Figure 1B). Participants were required to recall at least 40% of the associations to be included in the study.

### Targeted memory reactivation

Participants in the Reactivation group received incomplete auditory reminders during NREM sleep, following our previous protocol (Forcato et al., 2020). Each reminder consisted of a previously learned environmental sound followed by the first syllable of its associated word. The environmental sound was presented for approximately 3 s and continued while the first syllable was presented auditorily (1.5 s; approximately 45 dB). Consecutive reminders were separated by at least 7 s (Figure 1C). The 30 associations were presented in pseudorandom order and, when sleep duration permitted, the entire set was presented twice.

Cueing began after at least 10 min of stable SWS. When participants did not reach SWS, cues were presented during stable S2 containing frequent slow waves. Stimulation was suspended at the first sign of arousal, awakening, or transition to another sleep stage and resumed only after stable NREM sleep had been re-established. Participants received a mean of 46.92 ± 5.44 reminders.

Participants in the No-Reactivation group underwent the same nap procedure but received no memory-related cues. Continuous white noise was presented throughout the nap in both groups.

### Memory test

Memory was tested after the nap, approximately 24 h after training. Participants heard each environmental sound and verbally reported the associated word. No syllabic cue or corrective feedback was provided (Figure 1D).

The primary behavioral outcome was memory change, calculated as the number of correctly recalled words at test minus the number recalled at training. Values closer to zero therefore reflected better retention, whereas increasingly negative values reflected greater forgetting.

### Polysomnography and sleep scoring

Sleep was recorded using electroencephalography (EEG), electrooculography (EOG), and electromyography (EMG). EEG was recorded from F3, F4, C3, C4, P3, and P4 according to the international 10-20 system and referenced to the averaged mastoids. Signals were acquired with BrainAmp amplifiers (Brain Products GmbH, Munich, Germany), sampled at 250 Hz, and filtered between 0.16 and 35 Hz.

Sleep was scored offline in 30-s epochs according to Rechtschaffen and Kales criteria (1968) as wake, stage 1 (S1), stage 2 (S2), slow-wave sleep (SWS; stages 3 and 4), or rapid eye movement (REM) sleep. Total sleep time, time spent in each sleep stage, and stage percentages were calculated for each participant.

### Sleep EEG analyses

Spectral power was calculated for NREM sleep (S2 + SWS) using Fast Fourier Transformations and averaged across electrodes. Frequency bands were defined as slow oscillation (0.5–1 Hz), delta (1–4 Hz), theta (4–8 Hz), slow spindle (9–12 Hz), and fast spindle (12–15 Hz). Slow oscillations, fast sleep spindles, and slow oscillation–fast spindle coupling were detected and quantified using SpiSOP (Spindles, Slow Oscillations and Power Spectral Density; RRID:SCR_015673), an open-source MATLAB toolbox based on FieldTrip (RRID:SCR_004849) (Oostenveld et al., 2011; Klinzing et al., 2016; Rudzik et al., 2018; Cha et al., 2020). Slow oscillations were detected between 0.5 and 1.25 Hz and fast spindles between 12 and 15 Hz. Event density was expressed as the number of detected events per minute of NREM sleep. Slow oscillation–fast spindle coupling was defined on the basis of temporal co-occurrence between detected slow oscillations and fast spindles, and coupling density was calculated as the number of coupled events per minute of NREM sleep.

### Cue-related EEG analyses

EEG activity surrounding auditory cue presentation was analyzed in 8-s epochs extending from −2 to +6 s relative to sound onset. Cue onset was automatically marked in the Reactivation group. For the No-Reactivation group, surrogate markers were placed post hoc to approximate the number and temporal distribution of events in the Reactivation group. These markers were restricted to stable NREM sleep and excluded sleep-stage transitions and artifact-contaminated periods. Epochs containing arousals, awakenings, or artifacts were excluded. Time–frequency analyses were performed on the averaged central electrodes, with spectral power normalized to the −2 to 0 s prestimulus baseline.

### Statistical analysis

Analyses were performed using SPSS (IBM Corp.) and Python, with statistical significance set at α = 0.05. Training performance and memory change were compared between groups using independent-samples *t*-tests. Sleep architecture, spectral power, slow oscillation density and count, fast spindle density and count, and slow oscillation–spindle coupling were compared between groups using independent-samples *t*-tests or Mann–Whitney *U* tests when parametric assumptions were not met. Associations between sleep measures and absolute memory change were assessed using Pearson correlations. For the cue-related time–frequency analysis, groups were compared at each time–frequency point using Mann–Whitney *U* tests followed by cluster-based permutation correction with 1,000 permutations implemented in MNE-Python.

To assess whether the group difference in memory change remained after accounting for NREM sleep physiology, ANCOVA models were conducted with memory change as the dependent variable and Group as a fixed factor. Separate models included percentage of NREM sleep, NREM slow oscillation power, or slow oscillation count as covariates. An additional model included slow oscillation density and fast spindle density simultaneously as covariates. As a complementary sensitivity analysis, the analyses were repeated in subsets of participants whose values for the relevant NREM measures fell within the range shared by both groups. The overlapping range was defined as the intersection of the minimum and maximum values observed in the Reactivation and No-Reactivation groups, and memory change was compared between groups within each restricted subset.

## Results

### Memory performance

There were no significant differences in training performance between the Reactivation and No-Reactivation groups (Reactivation: 20.69 ± 1.35; No-Reactivation: 20.69 ± 1.15; *p* = 1.00). In contrast, memory change differed significantly between groups, with less forgetting in the Reactivation group than in the No-Reactivation group (−2.62 ± 0.29 vs. −4.92 ± 0.51; U = 28.0, p = 0.003, r = 0.67; Figure 2A).

**Figure 2.**
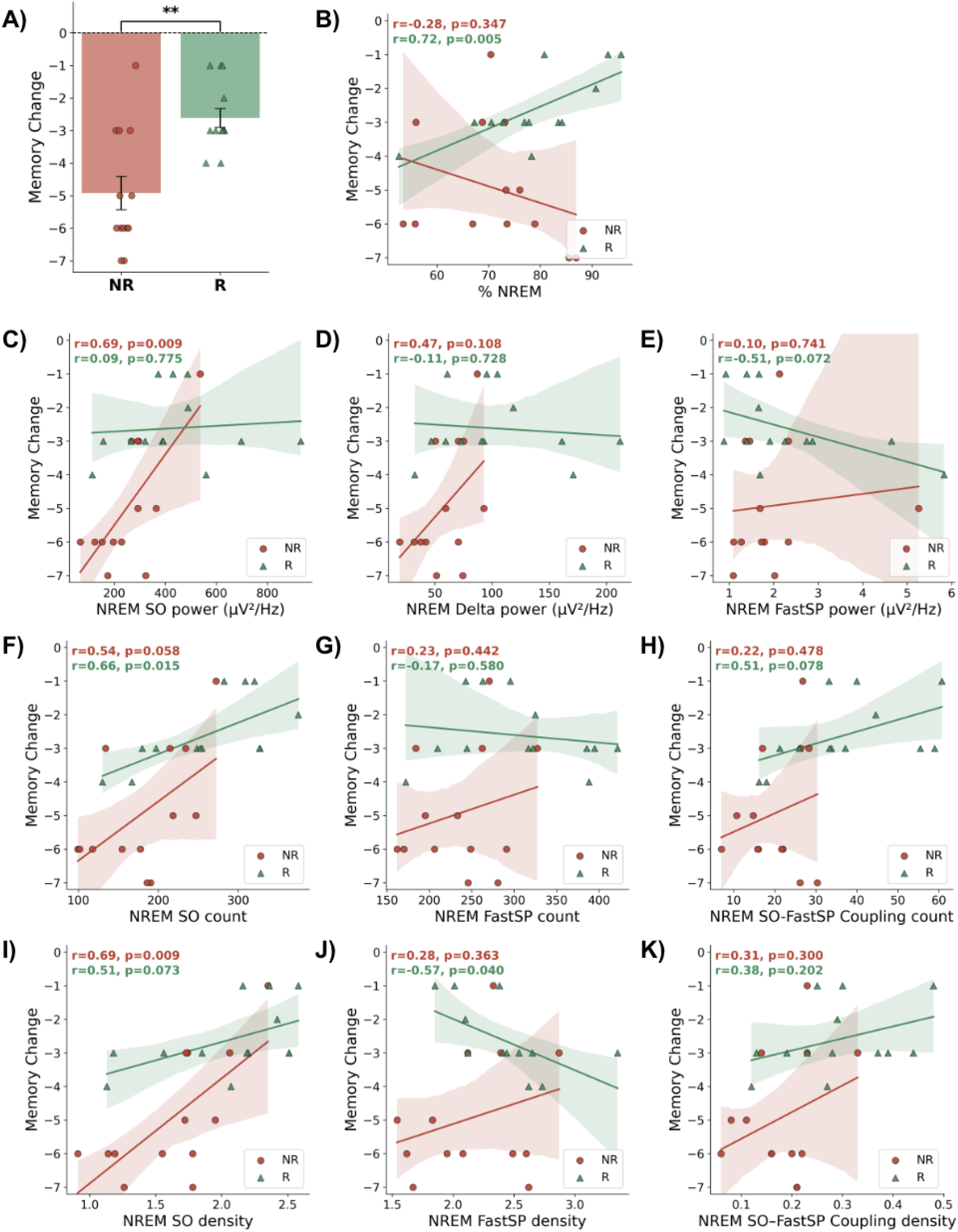
Memory change and NREM sleep measures in the Reactivation and No-Reactivation groups. **(A)** Comparison of memory change between the NR (red, n = 13) and R (green, n = 13) groups. Bars represent the mean ± SEM, and individual subjects are shown (circles: NR; triangles: R). Statistical significance is indicated by asterisks (**, p < 0.01). **(B-K)** Correlations between memory change and NREM sleep measures, including the percentage of NREM sleep, spectral power in the SO, delta, and fast spindle bands, the number of detected SOs, fast spindles, and SO-fast spindle couplings, as well as the density of these events. Pearson correlation coefficients (r) and corresponding p-values are shown separately for the NR (red) and R (green) groups. Solid lines represent the least-squares linear regression for each group, and shaded areas indicate the 95% confidence intervals.

### Sleep architecture and NREM sleep physiology

No significant differences were found between groups in total sleep time, or in the percentage of total sleep time spent in each stage (all *p* > 0.066; Table 1). Across NREM sleep, the Reactivation group showed higher slow oscillation (0.5–1 Hz, *p* = 0.017) and delta power (1–4 Hz, *p* = 0.012) than the No-Reactivation group, whereas theta (4-8 Hz) and spindle-band (12-15 Hz) power did not differ between groups (all *p* > 0.288) (Table 1).

**Table 1.** Comparison of sleep architecture, NREM spectral power density, and NREM event measures between the NR (control) and R (experimental) groups. Values are reported as mean ± SEM. Group comparisons were performed using independent-samples Student’s t-test when assumptions of normality and homogeneity of variances were met, and the Mann-Whitney U test when normality assumptions were violated. Cohen’s d is reported as the effect size for t-tests and rank-biserial correlation (r) for Mann-Whitney tests. Significant differences are highlighted in bold (p < 0.05).

|  | NR group (n=13) | R group (n=13) | Statistics |
| --- | --- | --- | --- |
| **Total sleep time (min)** | 82.50 ± 2.63 | 83.12 ± 4.25 | t(24)=-0.12, p=0.903, d=-0.05 |
| **Percentage of total sleep time spent in each stage** |  |  |  |
| **% W** | 2.81 ± 1.06 | 3.80 ± 1.75 | U = 84.0, p = 1.000, r = 0.01 |
| **% S1** | 19.41 ± 2.59 | 13.47 ± 2.18 | t(24) = 1.75, p = 0.092, d = 0.69 |
| **% S2** | 55.38 ± 3.01 | 54.25 ± 4.06 | t(24) = 0.22, p = 0.825, d = 0.09 |
| **% SWS** | 15.42 ± 2.04 | 24.58 ± 4.29 | t(24) = -1.93, p = 0.066, d = -0.76 |
| **% REM** | 7.12 ± 1.61 | 3.95 ± 1.51 | U = 114.0, p = 0.126, r = -0.35 |
| **% NREM** | 70.66 ± 2.95 | 78.77 ± 3.21 | t(24) = -1.86, p = 0.075, d = -0.73 |
| **NREM Power Spectral Density** |  |  |  |
| **SO (μV<sup>2</sup>/Hz)** | 254.64 ± 33.46 | 431.08 ± 60.30 | **t(24) = -2.56, p = 0.017, d = -1.00** |
| **Delta (μV<sup>2</sup>/Hz)** | 58.61 ± 6.14 | 101.40 ± 14.58 | **t(24) = -2.71, p = 0.012, d = -1.06** |
| **Theta (μV<sup>2</sup>/Hz)** | 7.43 ± 0.53 | 8.69 ± 1.03 | t(24) = -1.09, p = 0.288, d = -0.43 |
| **Fast spindle (μV<sup>2</sup>/Hz)** | 1.96 ± 0.30 | 2.30 ± 0.40 | U = 78.5, p = 0.778, r = 0.07 |
| **NREM event detection** |  |  |  |
| **SO count** | 180.60 ± 15.63 | 259.22 ± 20.37 | **t(24) = -3.06, p = 0.005, d = -1.20** |
| **Fast spindle count** | 236.61 ± 14.02 | 306.28 ± 21.52 | **t(24) = -2.71, p = 0.012, d = -1.06** |
| **SO-FastSP Coupling count** | 20.24 ± 2.01 | 36.83 ± 4.13 | **t(24) = -3.61, p = 0.001, d = -1.42** |
| **δSO (events/epoch)** | 1.63 ± 0.11 | 2.06 ± 0.14 | **U = 38.0, p = 0.018, r = 0.55** |
| **δFastSP (events/epoch)** | 2.16 ± 0.12 | 2.41 ± 0.11 | t(24) = -1.54, p = 0.137, d = -0.60 |
| **δSO-FastSP Coupling (events/epoch)** | 0.18 ± 0.02 | 0.29 ± 0.03 | **t(24) = -2.95, p = 0.007, d = -1.16** |

We also analyzed slow oscillations, fast spindles, and slow oscillation-spindle coupling as discrete events. Across NREM sleep, the Reactivation group showed a higher slow oscillation count (*p* = 0.005) and density (*p* = 0.018), as well as a higher fast spindle count (*p* = 0.012). Slow oscillation-fast spindle coupling was also higher in the Reactivation group, both in count (*p* = 0.001) and density (*p* = 0.007). Fast spindle density did not differ between groups (*p* = 0.137; Table 1).

### Associations between sleep architecture, NREM physiology and memory

Correlations between memory change and NREM sleep measures were examined separately for the Reactivation and No-Reactivation groups (Figure 2B-K). In the Reactivation group, memory change was positively associated with the percentage of NREM sleep (*r* = 0.72, *p* = 0.005) and with slow oscillation count (*r* = 0.66, *p* = 0.015), indicating that greater NREM sleep and a higher number of slow oscillations were associated with less forgetting. Fast spindle density showed a negative association with memory change in this group (*r* = −0.57, *p* = 0.040), with higher spindle density associated with greater forgetting.

In the No-Reactivation group, memory change was positively associated with NREM slow oscillation power (*r* = 0.69, *p* = 0.009) and slow oscillation density (*r* = 0.69, *p* = 0.009), indicating less forgetting in participants with greater slow oscillatory activity. No other correlations between memory change and the NREM sleep measures shown in Figure 2B-K reached statistical significance.

### Memory effects after accounting for sleep physiology

Because several NREM sleep measures differed between groups or were associated with memory retention, we examined whether these variables accounted for the behavioral effect of reactivation.

The effect of Group on memory change remained significant after controlling separately for the percentage of NREM sleep (*F*(1,23) = 11.86, *p* = 0.002, partial η^2^ = 0.340), NREM slow oscillation power (*F*(1,23) = 8.02, *p* = 0.009, partial η^2^ = 0.259), and slow oscillation count (*F*(1,23) = 5.00, *p* = 0.035, partial η^2^ = 0.179). The group effect also remained significant in a model including both slow oscillation density and fast spindle density as covariates (*F*(1,22) = 5.07, *p* = 0.035, partial η^2^ = 0.187; Table 2). Within these models, slow oscillation count (*F*(1,23) = 9.54, *p* = 0.005, partial η^2^ = 0.293) and slow oscillation density (*F*(1,22) = 11.60, *p* = 0.003, partial η^2^ = 0.345) were significantly associated with memory change, whereas percentage of NREM sleep, slow oscillation power, and fast spindle density were not significant covariates (all *p* ≥ 0.109).

**Table 2.** Analysis of covariance (ANCOVA) examining the effect of group on memory change after adjusting for sleep-related covariates. Results of analyses of covariance (ANCOVA) examining whether the difference in memory change between the NR and R groups remained after adjusting for sleep variables. Four separate ANCOVA models were fitted, including either %NREM sleep, NREM slow oscillation (SO) power, SO count, or both SO density and fast spindle density as covariates. The dependent variable was memory change, and group (NR vs. R) was included as a fixed factor in all models. F statistics, associated p-values, and partial eta squared (partial η^2^) are reported for both the group effect and each covariate.

| ANCOVA model | Effect | F(df1, df2) | p | Partial $\eta^2$ |
| --- | --- | --- | --- | --- |
| %NREM | Group | **11.86 (1,23)** | **0.002** | **0.340** |
|  | %NREM | 0.21 (1,23) | 0.652 | 0.009 |
| NREM SO power | Group | **8.02 (1,23)** | **0.009** | **0.259** |
|  | NREM SO power | 2.78 (1,23) | 0.109 | 0.108 |
| NREM SO count | Group | **5.00 (1,23)** | **0.035** | **0.179** |
|  | SO count | 9.54 (1,23) | 0.005 | 0.293 |
| SO density + Fast spindle density | Group | 5.07 (1,22) | 0.035 | 0.187 |
|  | SO density | 11.60 (1,22) | 0.003 | 0.345 |
|  | Fast spindle density | 0.45 (1,22) | 0.508 | 0.020 |

As a complementary sensitivity analysis, we restricted the sample to participants whose NREM slow oscillation power fell within the range shared by both groups. The Reactivation group continued to show significantly less forgetting than the No-Reactivation group (R: −2.40 ± 0.34; NR: −4.83 ± 0.55; *U* = 19.0, *p* = 0.006, *r* = 0.68; Figure 3). Importantly, the between-group difference in memory change remained significant in all sensitivity analyses using overlapping ranges of the remaining NREM measures (Supplementary Figure 1).

**Figure 3.**
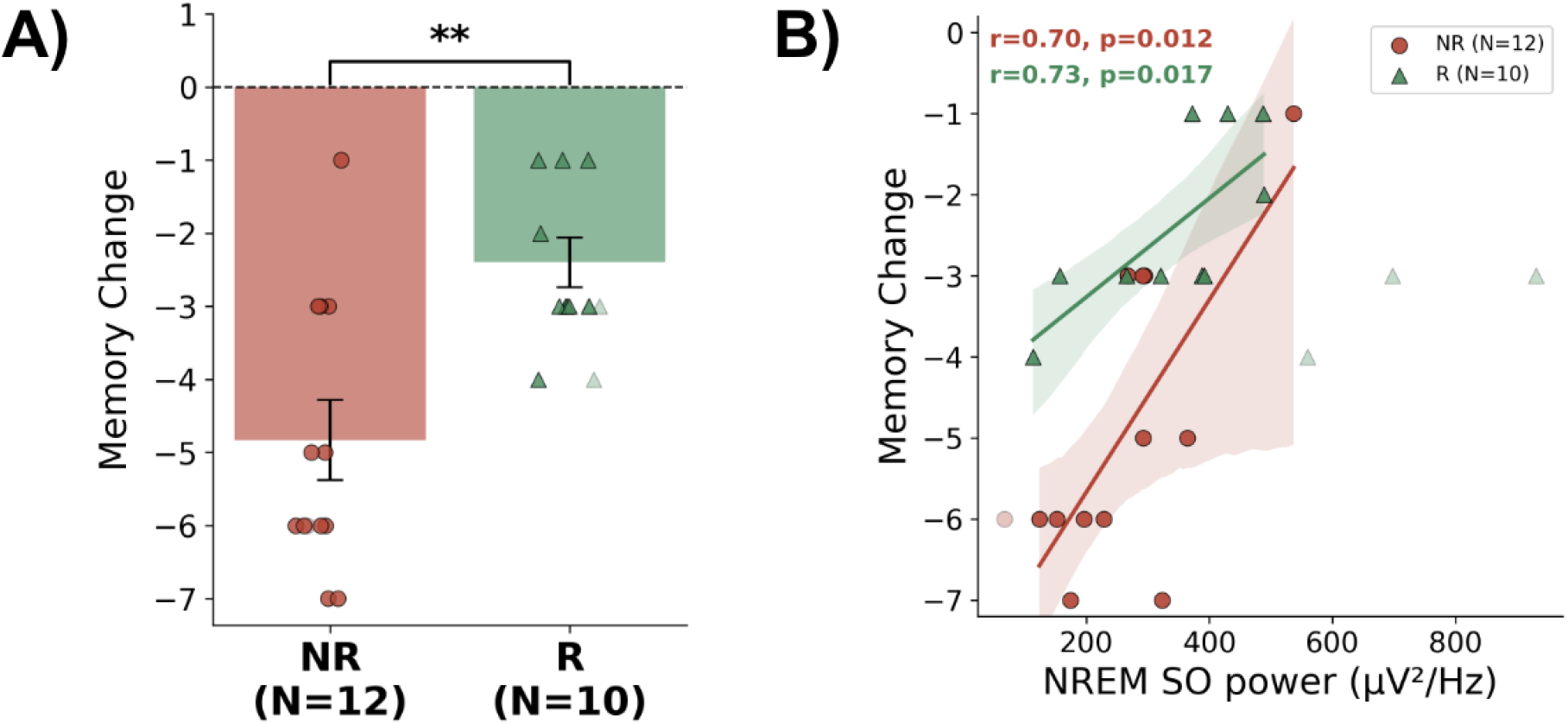
Sensitivity analysis after restricting NREM sleep measures to overlapping ranges between groups. Memory change after restricting the sample to the overlapping range of NREM slow oscillation power. Comparison of memory change between the NR (n = 12) and R (n = 10) groups after restricting the sample to participants with NREM slow oscillation power values within the range shared by both groups. Bars represent mean ± SEM, and individual participants are shown (circles: NR; triangles: R). Participants outside the overlapping range are shown with transparency and were excluded from the statistical analysis. p < 0.01.

### Cue-related electrophysiological responses

Time–frequency analysis revealed transient changes in spectral power following presentation of the incomplete auditory reminders (Figure 4). Compared with surrogate NREM periods in the No-Reactivation group, significant between-group differences were observed in the slow oscillation, delta, theta, and fast spindle ranges following environmental sound onset (all cluster *p* < 0.033). Following syllabic cue onset, significant differences were observed in the slow oscillation, delta, and theta ranges (all cluster *p* < 0.004), whereas no significant difference was found in the fast spindle range (*p* = 0.89).

**Figure 4.**
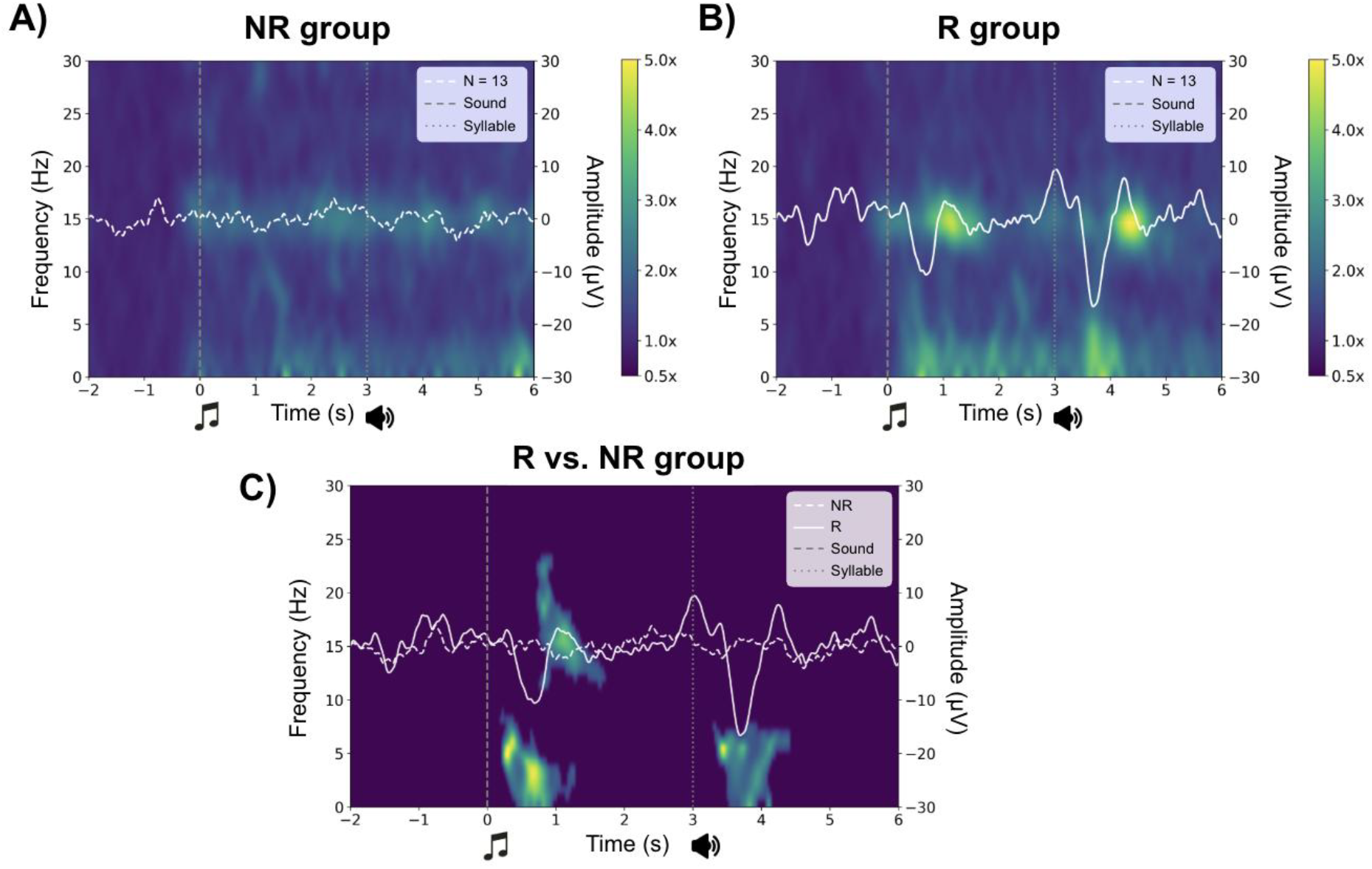
Cue-related time–frequency activity during NREM sleep. (A) Grand-average baseline-normalized spectrogram for the No-Reactivation group (NR; N = 13), time-locked to surrogate markers placed during stable NREM sleep. (B) Grand-average baseline-normalized spectrogram for the Reactivation group (R; N = 13), time-locked to presentation of the auditory reminders. (C) Time– frequency map of significant between-group differences identified using a non-parametric cluster-based permutation analysis (1,000 permutations, two-tailed). Mann–Whitney *U* tests were performed at each time–frequency point, and clusters were formed from adjacent points exceeding the initial significance threshold (*p* < 0.05) before cluster-level correction. In all panels, overlaid traces represent the grand-average event-related potential (ERP). The dashed and dotted vertical lines indicate environmental sound onset and syllabic cue onset, respectively. Spectral power was normalized to the −2 to 0 s prestimulus baseline.

### Correlations between spectral power clustering and memory performance

To examine whether the magnitude of the cue-related electrophysiological responses was associated with memory retention, we correlated memory change with spectral power extracted from each of the three significant time–frequency clusters in the Reactivation group. No significant associations were observed. Memory change was not related to activity in the first sound-related cluster, comprising slow oscillation, delta, and theta activity (*r* = −0.08, *p* = 0.783), or in the second sound-related cluster, corresponding to fast spindle activity (*r* = 0.06, *p* = 0.839). Likewise, activity in the cluster following the syllabic cue, comprising slow oscillation, delta, and theta activity, was not significantly associated with memory change (*r* = −0.40, *p* = 0.181).

## Discussion

Here we showed that TMR during NREM sleep reduced forgetting of sound–word associations learned 24 h earlier. Participants who received reminders during the nap retained more associations than participants who slept without memory-related cues, despite no significant differences in training performance or sleep macroarchitecture. Most TMR studies have presented cues during the first sleep period after learning, when post-encoding consolidation is still prominent (Rasch et al., 2007; Rudoy et al., 2009; Antony et al., 2012; Schreiner & Rasch, 2015). The present results extend this work by showing that sleep cueing can influence declarative memory after a 24-h consolidation interval. To our knowledge, this provides the first evidence in humans that TMR can benefit declarative memories reactivated at this later time point.

Evidence that sleep reactivation can affect memories beyond the immediate post-learning period has previously come from animal work. Rolls et al. (2013) re-exposed mice to a conditioned odor during sleep 24 h after fear conditioning and found enhanced subsequent memory expression. Blocking protein synthesis in the basolateral amygdala altered this effect, indicating that reactivation during sleep can recruit plasticity mechanisms after the initial learning period. The present findings extend this observation to human declarative memory, although they do not establish that the underlying mechanisms are the same.

The behavioral effect was accompanied by marked differences in NREM sleep physiology. Across the nap, the Reactivation group showed greater slow oscillation and delta power, higher slow oscillation density and count, greater fast spindle count, and higher slow oscillation–spindle coupling density and count. Several of these measures were also associated with memory retention. One possibility is that repeated auditory stimulation itself contributed to the increase in slow-frequency activity. Auditory stimuli presented during NREM sleep can elicit slow-wave responses, and repeated presentation of reminders may therefore have increased the occurrence of slow oscillations beyond the immediate cue-related epochs. In the present protocol, the complete set of reminders was presented twice, providing greater cue exposure than in our previous study using the same task (Forcato et al., 2020). This greater stimulation load may have contributed to the differences in slow oscillatory activity observed across the nap.

An increase in slow oscillations would also provide more opportunities for temporal co-occurrence with sleep spindles (Mölle et al., 2011; Staresina et al., 2015). Thus, the higher number and density of coupled events observed in the Reactivation group may partly reflect the greater availability of slow oscillations rather than a change in coupling alone. Nevertheless, the global physiological differences do not provide a simple explanation for the behavioral effect. The group difference in memory remained significant after statistical adjustment for the NREM measures and when memory performance was compared in participants with overlapping ranges of these variables. The memory benefit therefore cannot be reduced to the Reactivation group simply exhibiting more slow oscillations, spindles, or coupled events. These analyses should not be interpreted as showing that sleep oscillations were mechanistically unrelated to the memory effect. The physiological measures were obtained after group assignment and may themselves have been influenced by the reminder cues. They could therefore form part of the processes initiated by reactivation rather than acting as independent covariates. What the analyses indicate is more limited: differences in the amount of the measured NREM activity did not statistically account for the full difference in memory performance between groups.

When we examined cue-locked EEG activity during TMR, auditory reminders elicited significant time-frequency responses in the slow oscillation, delta, theta, and fast-spindle ranges. However, power extracted from the significant clusters was not associated with individual differences in memory change. Thus, although the reminders elicited robust electrophysiological responses, the magnitude of these immediate responses was not directly related to the subsequent memory benefit. One possibility is that a memory reminder initiates processing that extends beyond the few seconds surrounding cue presentation. The auditory cue may reactivate the corresponding memory representation and trigger subsequent plastic processes that continue after the stimulus has ended. Evidence from wakefulness supports this possibility. Bavassi et al. (2026) identified changes in spectral activity and functional network organization during the resting period following memory retrieval, indicating that the neural state induced by a reminder can persist beyond the retrieval event itself. Although that study was conducted during wakefulness and used a different memory paradigm, it provides a useful conceptual parallel: during sleep, memory reactivation may similarly initiate processes that continue during subsequent NREM sleep rather than being confined to the immediate cue-evoked response.

This interpretation is also compatible with our previous findings using the same sound-word paradigm. In Forcato et al. (2020), electrophysiological responses elicited during TMR did not show a consistent relationship with subsequent memory performance, despite the behavioral effects of memory cueing. A similar dissociation was observed here: reminders elicited significant EEG responses, but the magnitude of these responses was not associated with memory change. Together, these findings suggest that the immediate electrophysiological response to a reminder may not capture the full mnemonic processing initiated by that cue. Reactivation may instead trigger changes that continue beyond stimulation and unfold during subsequent sleep.

At the same time, the Reactivation group showed broader differences in NREM physiology across the nap, including greater slow oscillation and delta power and differences in slow oscillations, spindles, and their coupling. Repeated auditory stimulation may itself have contributed to some of these global changes, particularly the increase in slow oscillatory activity. However, the behavioral difference between groups remained after accounting for these NREM measures, indicating that the memory benefit cannot be reduced simply to the Reactivation group exhibiting greater slow oscillatory or spindle-related activity.

The use of incomplete reminders may be particularly relevant in this context. Incomplete cues provide only part of the expected information and may therefore promote retrieval while leaving the associated representation unresolved. During wakefulness, reminders that generate a mismatch between expected and presented information can render established memories susceptible to further modification (Forcato et al., 2007, 2009, 2011; Fernández et al., 2016; Sinclair & Barense, 2018). In the present study, incomplete reminders again reduced subsequent forgetting, now when cueing occurred 24 h after learning. However, because no complete-reminder condition was included, the current experiment cannot determine whether incomplete information was necessary for the effect or whether complete reminders would have produced a similar benefit.

The present findings can also be considered in the context of memory reactivation during wakefulness. Retrieval of an established memory can make it susceptible to subsequent modification (Nader et al., 2000; Sara, 2000; Dudai, 2012), and in human declarative memory, repeated reactivation with incomplete reminders can produce strengthening that becomes evident after a delay (Forcato et al., 2011, 2013). The present results suggest that a similar principle may operate during sleep. Rather than producing an effect confined to the moment of cue presentation, a reminder may initiate a period of post-reactivation processing during which the memory remains susceptible to further modification. In this view, the behavioral benefit observed after the nap may reflect processes initiated by the reminder that continue during subsequent NREM sleep. Whether these post-reactivation processes overlap with mechanisms described for memory reconsolidation during wakefulness remains to be determined.

Several limitations are relevant to this interpretation. First, the final sample was relatively small, with 24 of the 50 recruited participants excluded. Most exclusions were related to the difficulty of obtaining sufficient stable NREM sleep and an adequate number of reminder presentations, rather than to technical problems alone. This is particularly relevant in the present design, in which TMR was performed during a daytime nap. Sleep propensity and stability are shaped by both homeostatic and circadian processes, and homeostatic sleep pressure is generally lower during the daytime than near the beginning of nocturnal sleep (Borbély, 2016). Maintaining stable NREM sleep while presenting auditory reminders may therefore be especially challenging in a short daytime nap. Auditory cues must be sufficiently salient to reactivate the associated memory while remaining subtle enough to avoid arousals or awakenings (Oudiette & Paller, 2013). This balance may be particularly important for verbal material, as words and speech are behaviorally meaningful stimuli, and auditory-induced sleep disturbances have been shown to reduce the effectiveness of TMR (Göldi & Rasch, 2019). By contrast, olfactory stimulation has a substantially lower capacity to induce awakening during sleep (Carskadon & Herz, 2004). Thus, the relatively high exclusion rate may partly reflect the combination of a daytime nap and the use of salient auditory reminders, and future studies should examine whether individualized stimulus intensity or longer sleep opportunities can improve retention of participants without compromising effective cueing. Second, the No-Reactivation group did not receive discrete auditory stimuli unrelated to the learned material. Continuous white noise was present in both groups, but this does not control for the physiological effects of individual sounds during sleep. Consequently, some of the cue-related and whole-nap physiological differences may reflect auditory stimulation itself rather than mnemonic reactivation. A future design including acoustically matched, non-associated cues, or cued and uncued memories within the same participant, would help separate these components. Third, a 24-h delay places reactivation beyond the immediate post-encoding period but does not imply that memory consolidation is complete. The findings are therefore better interpreted as evidence for modification after an initial consolidation interval rather than after a definitively completed consolidation process. Finally, memory was tested shortly after the nap. Whether the reduction in forgetting persists over subsequent days remains to be established.

## Conclusion

Targeted memory reactivation during NREM sleep reduced forgetting of declarative memories learned 24 h earlier. Reactivation was accompanied by both immediate cue-related responses and broader changes in NREM physiology. Although repeated auditory stimulation may have contributed to the increase in slow oscillatory activity across the nap, differences in slow oscillations, spindles, and their coupling did not statistically account for the full behavioral effect. This dissociation suggests that the mnemonic consequences of a reminder may not be confined to the few seconds surrounding its presentation. Instead, cueing may initiate memory processing that continues during subsequent sleep. These findings broaden the temporal window over which sleep reactivation can influence memory and raise the possibility that memories reactivated after an initial consolidation interval remain susceptible to further plastic modification.

## Author contribution

Malen D. Moyano: Conceptualization; methodology; investigation; writing original draft; writing - review and editing; data curation. Lucila Capurro: investigation; visualization; formal analysis; writing original draft; writing - review and editing. María C. González: investigation; visualization; writing - review and editing. Luis I. Brusco: Review and editing; resources. Cecilia Forcato: Conceptualization, methodology; writing original draft; writing - review and editing; resources; project administration; supervision.

## Data availability statement

All behavioral and electrophysiological data generated and analyzed during this study will be made publicly available in an open-access repository upon acceptance of the manuscript. Repository details and persistent identifiers (DOI) will be provided before publication to ensure unrestricted access to the data supporting the findings of this study.

## Institutional Review Board Statement

The study was conducted in accordance with the Declaration of Helsinki and was approved by the Biomedical Research Ethics Committee of the Instituto Alberto C. Taquini, University of Buenos Aires. Written informed consent was obtained from all participants included in the study.

## Use of Artificial Intelligence

The authors used ChatGPT (OpenAI) to assist with translation and English language editing of the manuscript, as the authors’ native language is Spanish. No AI tools were used for data analysis, interpretation of results, or generation of scientific conclusions. The authors reviewed and edited all AI-assisted text and figure-related suggestions and take full responsibility for the final content of the manuscript.

