## Supplementary Information for "TARGETED MEMORY REACTIVATION DURING NON-RAPID EYE MOVEMENT SLEEP STRENGTHENS CONSOLIDATED DECLARATIVE MEMORIES"

### Supplementary Results

To further examine whether the difference in memory change between groups depended on between-group differences in NREM sleep physiology, analyses were repeated after restricting the samples to participants whose values fell within the range shared by the Reactivation and No-Reactivation groups for each NREM measure (Supplementary Figure 1). After restriction based on the percentage of NREM sleep, the remaining sample comprised 10 participants in the Reactivation group and 13 in the No-Reactivation group. The Reactivation group showed significantly less forgetting than the No-Reactivation group (R: −3.00 ± 0.26; NR: −4.92 ± 0.51; $U$ = 26.0, $p$ = 0.012, $r$ = 0.60; Supplementary Figure 1A). Within this restricted sample, memory change was not significantly associated with percentage of NREM sleep in either group (NR: $r$ = −0.28, $p$ = 0.347; R: $r$ = 0.44, $p$ = 0.199; Supplementary Figure 1B). Restriction based on slow oscillation count resulted in a smaller sample, comprising 8 participants in the Reactivation group and 10 in the No-Reactivation group. The between-group difference in memory change did not reach statistical significance (R: −3.00 ± 0.33; NR:

−4.60 ± 0.64; *U* = 21.0, *p* = 0.085, *r* = 0.47; Supplementary Figure 1C). Within this restricted sample, slow oscillation count was positively associated with memory change in the Reactivation group (*r* = 0.78, *p* = 0.023), whereas the association was not significant in the No-Reactivation group (*r* = 0.47, *p* = 0.175; Supplementary Figure 1D). After restriction based on slow oscillation density, 9 participants remained in the Reactivation group and 12 in the No-Reactivation group. The Reactivation group continued to show significantly less forgetting (R: −2.78 ± 0.36; NR: −4.83 ± 0.55; *U* = 21.5, *p* = 0.019, *r* = 0.60; Supplementary Figure 1E). Slow oscillation density was positively associated with memory change in the No-Reactivation group (*r* = 0.71, *p* = 0.009), but not in the Reactivation group (*r* = 0.53, *p* = 0.143; Supplementary Figure 1F). Finally, restriction based on fast spindle density resulted in 12 participants in the Reactivation group and 9 in the No-Reactivation group. The Reactivation group again showed significantly less forgetting than the No-Reactivation group (R: −2.58 ± 0.31; NR: −4.78 ± 0.66; *U* = 20.5, *p* = 0.015, *r* = 0.62; Supplementary Figure 1G). Fast spindle density was negatively associated with memory change in the Reactivation group (*r* = −0.72, *p* = 0.009), whereas no significant association was observed in the No-Reactivation group (*r* = −0.12, *p* = 0.754; Supplementary Figure 1H). Overall, the between-group difference in memory change remained significant after restriction based on percentage of NREM sleep, slow oscillation density, and fast spindle density. The only sensitivity analysis in which the group difference did not reach statistical significance was that based on slow oscillation count, which also resulted in the smallest remaining Reactivation sample (*n* = 8).

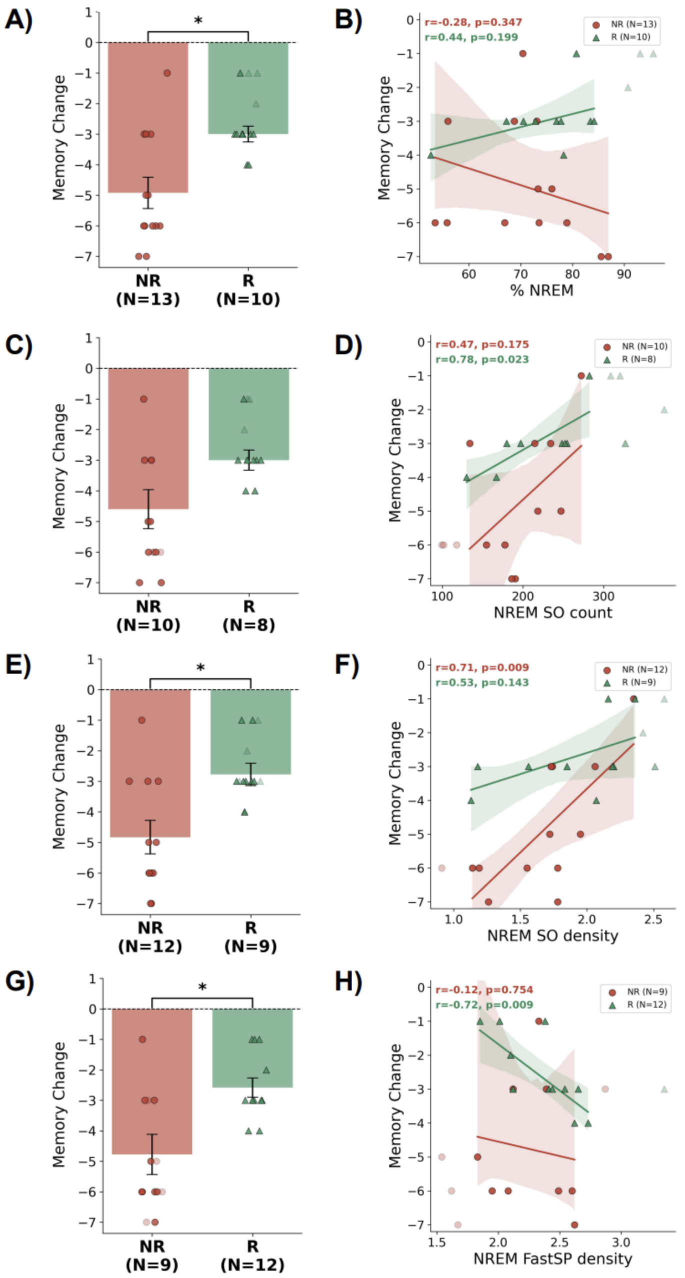


**Supplementary Figure 1. Sensitivity analysis of the relationship between memory change and additional NREM sleep measures.** Analyses were repeated after restricting each NREM measure to the range of values shared by the No-Reactivation (NR) and Reactivation (R) groups, defined as the intersection of the minimum and maximum values observed in both groups. Participants outside the overlapping range are shown with transparency and were excluded from the corresponding statistical analysis. **(A, B)** Analyses restricted according to percentage of NREM sleep. **(C, D)** Analyses restricted according to NREM slow oscillation (SO) count. **(E, F)** Analyses restricted according to NREM SO density. **(G, H)** Analyses restricted according to NREM fast spindle (FastSP) density. Left panels (A, C, E, G) show memory change in the NR and R groups; bars represent mean ± SEM and individual participants are shown as circles (NR) and triangles (R). Group comparisons were performed using Mann–Whitney *U* tests. Right panels (B, D, F, H) show correlations between memory change and the corresponding NREM measure within the restricted samples. Pearson correlation coefficients (*r*) and *p*-values are shown separately for each group. Solid lines represent least-squares linear regressions and shaded areas indicate 95% confidence intervals. $p < 0.05$.
